# Assessing a natural gene drive for population control in house mice

**DOI:** 10.64898/2026.10.13.757620

**Authors:** Andri Manser, Anna K Lindholm

## Abstract

House mice are a major invasive pest with significant impact on island ecosystems. Genetic biocontrol via gene editing offers new avenues for population control. An alternative idea dating back over half a century is to harness the unmodified, naturally occurring genetic element called *t* haplotype for population suppression. The *t* haplotype causes sterility in *t/t* homozygote males, while *+/t* heterozygote males transmit the *t* to most of their offspring via meiotic drive. We use population genetic modelling to systematically explore whether this approach can suppress invasive populations through the accumulation of *t/t* sterile males. We find that *+/t* fertile releases combined with the sustained release of *t/t* sterile males can achieve eradication, but require prolonged interventions, with eradication taking around 25 years under our baseline scenario. We then show that release effort and eradication times may be reduced substantially if releases are combined with trapping, culling, or the release of hypercompetitive (*Adonis*) males. A direct comparison with *t*_CRISPR_, currently the main engineered genetic biocontrol system for house mice, shows that the approach requires greater release effort but achieves comparable eradication times. We provide an R shiny application for exploring model scenarios interactively. Our findings highlight both the potential and the limitations of exploiting this natural meiotic drive system for population control, and identify key gaps in knowledge of mouse reproductive biology and behaviour needed to fully assess its feasibility in the field.

---

> “I wonder if anyone has thought of controlling the mouse population of an area by flooding it with *t*-carriers.”
>
> George C. Williams in *Adaptation and Natural Selection* (1966), page 119 (1)

House mice (*Mus musculus*) are a major ecosystem pest species. On the backwash of human activity, these ephemeral, versatile, and rapidly reproducing rodents have colonised virtually every corner of our planet, including about 80% of islands that are particularly vulnerable to biodiversity loss (2). As a non-native invasive species, they have devastating impact on local biota through predation, competition or disease transmission (3; 4). A dramatic example is Gough Island, where house mice are estimated to have killed 2 million eggs and chicks of seabird species annually, some critically endangered (5). Efforts to control invasive house mouse populations date back over half a century (6). Traditional control methods such as anticoagulant poisons and trapping have been applied successfully, but only about half of the 116 eradication attempts recorded since 1971 have been successful (6). The toxicant approach further raises serious animal welfare concerns (anticoagulants kill animals through slow internal bleeding), kill non-target species (directly and indirectly via predation), and is difficult to apply in areas populated by humans due to its effects on health. Mice are also evolving resistance to toxins (7; 8).

Rapid advances in gene editing technology have opened up interesting new avenues to control pest species via genetic engineering (9). Gene drives, defined as stretches of DNA that are passed on to systematically more than 50% of progeny expected under Mendelian inheritance, can now be synthesised in the lab. Thanks to their Super-Mendelian mode of inheritance, gene drives, along with any additional linked gene material (cargo), are expected to rapidly propagate in target populations, even if the construct incurs costs to the organisms that carry them. The potential for spread despite cost makes them promising candidates to genetically modify or eradicate pest species (10; 11). A functional synthetic gene drive based on *CRISPR/Cas* has been presented for female house mice (12), though the development of potent gene drives appears to be complicated compared to insects (13). Additionally, modification of a naturally occurring driver in house mice called *t* haplotype (see below) has been suggested for mouse control, either by coupling the driver to a male sex determination locus (*Sry*), thus eradicating target populations due to a lack of males (14; 11; 15), or to a CRISPR-Cas cassette triggering a loss-of-function in a female fertility gene (*Prl* (16)). Despite the great potential, synthetic gene drive technology raises a host of challenges. Empirical and theoretical studies suggest that resistance to a driver may evolve on temporal scales that could render interventions ineffective (17). Perhaps more importantly, the release of genetically engineered constructs designed for rapid spread in and modification of natural populations raises serious ethical, ecological, and political concerns (18).

An idea dating back over 50 years is the use of the naturally occurring gene drive *t* haplotype to control invasive mouse populations (19; 20; 1). The *t* haplotype is a variant of mouse chromosome 17 that has segregated in the *Mus musculus* cluster for over 3 million years (21). Two characteristics make the *t* haplotype promising for pest control: First, several of 16 described naturally occurring variants cause sterility in *t/t* homozygote males (22). The release of *t/t* males could thus be utilised to hamper reproductive rates by increasing the frequency of unproductive matings. Similar approaches have seen considerable success in insect agricultural and disease pests (referred to as the sterile (male) insect technique, (23)). Second, the *t* haplotype offers an additional twist: it is a potent gene drive. In *+/t* heterozygote males, sperm development is manipulated such that most *+* sperm are rendered dysfunctional. As a result, *+/t* heterozygote carriers transmit the *t* to 90–99% of their progeny instead of the 50% expected under Mendelian inheritance (24). Thanks to this systematic transmission advantage, the driver may genetically propagate in populations despite the sterility cost imposed on *t/t* males. Additional to the direct release of sterile *t/t* males, the release of *+/t* heterozygote males may thus be used to raise sterility levels indirectly via genetic propagation. The idea that sterile *t* haplotypes may trigger population extinctions —both in the absence and presence of active releases— has been around since the 1960s (19; 20; 1), but has never been explored systematically.

Our aim here is to assess the general feasibility of sterile *t* driven population suppression. To this end, we model *t* release campaigns by tracking the abundance of the three genotypes (*+/+, +/t, t/t*, Fig. 1A) in a system of coupled differential equations. While spatial structure is expected to play an key role in the effectiveness of releases, our goal here was to build a conceptual model assuming a panmictic (well-mixed) population, allowing us to assess whether suppression is achievable *in principle* across a broad range of biological parameters and release scenarios. The po-tential of the sterile *t* for pest control is assessed in three steps. First, we examine a baseline scenario based on current knowledge where *t* carrying males are released into a target population. Second, we assess if campaigns can be refined if *t* releases are combined with trapping, previous culling, and if males are released that are particularly attractive to females. We further directly compare sterile *t* with *t*_CRISPR_, the recently developed, genetically modified *t* haplotype version that induces female infertility via CRISPR/Cas (16). Third, we perform systematic sensitivity analysis, assessing how release effort and eradication times depend on a number of biological processes.

**Figure 1.**
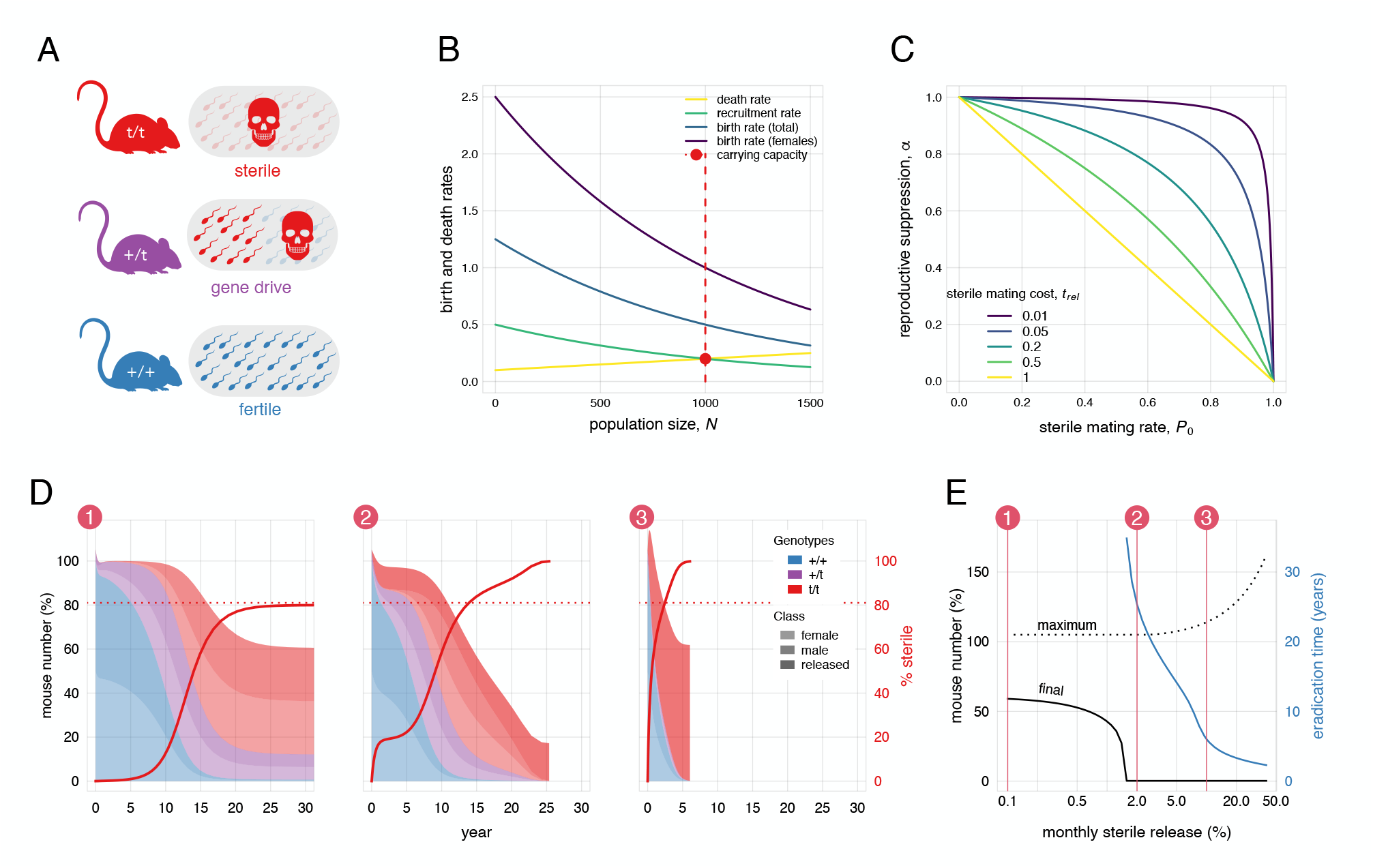
Using the sterile *t* for house mouse pest control. A. **A**. Male genotypes and phenotypes: *t/t* males are sterile with no functional sperm, *+/t* fertile males transmit (almost) only *t* sperm (gene drive), and *+/+* males reproduce normally. Females show no phenotypic differences. **B**. Demographic profile of the target population, characterized via density-dependent recruitment *b*(*N*) (combining birth rate and pup survival) and death rate *m*(*N*). Birth and death rates are equal at carrying capacity *K* (red dot). **C**. Reproductive suppression caused by sterile males (α) depends on *t/t* mating success (rate *P*_0_) and the reproductive cost of sterile matings, measured as the relative delay in a female’s reproductive schedule (*t*_rel_). **D**. Model dynamics for the baseline scenario (Tab. 1) under three release strategies: 1. Single 5% release of *+/t* heterozygotes. 2. Single 5% *+/t* and monthly 2% *t/t* release. 3. Single 5% *+/t* and 10% monthly *t/t* release. Shown the abundance of the three genotypes (*+/+* blue, *+/t* violet, *t/t* red) for both sexes (females light, males medium, released males dark), stacked cumulatively as proportions of carrying capacity (*K*). Red lines indicate the frequency of sterile *t/t* males (*y*_*t*/*t*_). The red dotted line shows the equilibrium prediction by Dunn (19). **E**. Population size at the end of the release campaign (black solid), maximum population size (black dotted), and eradication time (blue) for systematically varying levels of monthly *t/t* sterile male release rate (on a log-10 scale). The three scenarios shown in D are indicated by numbers and red vertical lines.

## Results

### Baseline model

The baseline model is based on current knowledge of house mouse biology and the *t* system (Table 1). Females produce 0.4 offspring per month, increasing exponentially to 2.5 as population size decreases (25), of which 40% survive into adulthood (26) (Fig. 1B). All adults die at a monthly rate of 20%, which drops to 10% as population size decreases (25). Females are polyandrous, either mating with one (monandry) or two (polyandry) males with a polyandry rate of 47%, which is at the up-per end of the spectrum reported in natural populations (27; 28; 29). *t* males (*+/t* and *t/t*) have the same ability to survive, establish territories and acquire matings as *+/+* males (30). Crucially, if a female mates with *t/t* sterile males only, she produces no offspring and experiences a delay of 19 days in her reproductive schedule (31) relative to the average interbirth interval of 67 days in wild house mice (32), which corresponds to a reproductive cost of 30% imposed by *t/t* sterile males (Fig. 1C). *+/t* fertile males transmit the *t* to 95% of their progeny (33; 34; 16), but only fertilise 11% of ova when in sperm competition against *+/+* males, corresponding to +/*t* male sperm competitiveness of 12.6% (35; 36; 37). Fertile males that compete with *t/t* sterile males will fertilise the entire litter.

**Table 1.**
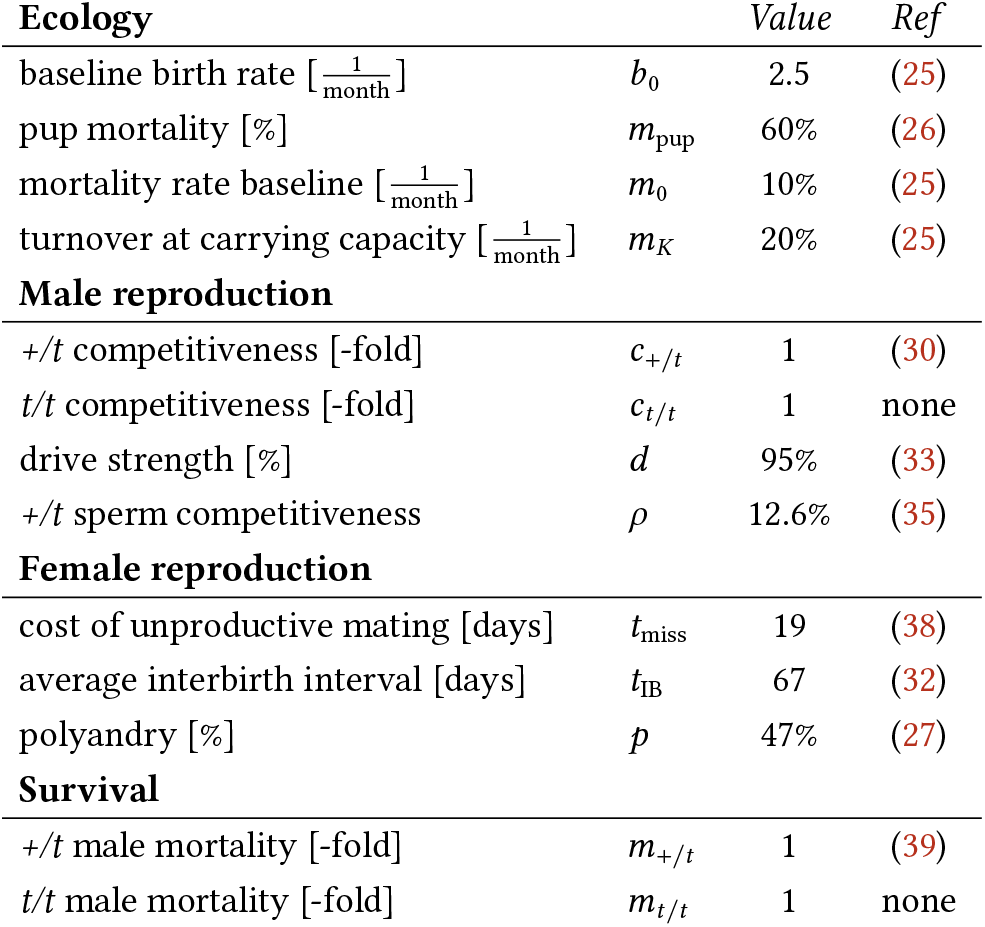
Model parameters and estimates of the baseline scenario.

### Genetic *t* spread

Figure 1D.1 shows the dynamics af-ter *+/t* fertile males are released once at 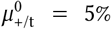 of carrying capacity. The *t* spreads genetically thanks to gene drive in *+/t* heterozygote males, but eventually slows as *t/t* steriles become common. After roughly 20 years, gene frequency stabilizes at a polymorphic equilibrium of 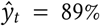 (corresponding to 80% *t/t* sterile males). This closely matches the analytical prediction by Dunn (19), who calculated that the opposing forces of gene drive promoting and male sterility opposing the *t* will balance at 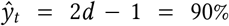. Polyandry and sperm competition do not appreciably suppress the *t* here due to negative frequency-dependence: when *t* is common, *+/t* males mostly compete with *+/t* and *t/t* rivals, to which they are equal and superior. Only low *t* frequency exposes *+/t* males to sperm competition against superior *+/+* males, which can create invasion thresholds (Fig. S3).

The stable presence of *t/t* sterile males hampers birth rates, reducing population size to 61% of carrying capacity, but sterile impact is not sufficient for population collapse. In Text S1, we analytically derive the % sterile males necessary for eradication, which is 95.2% for our baseline parameter settings. Direct eradication following a single release is only possible for a relatively narrow parameter range (Fig. S1–2).

### Adding steriles

We can combine *+/t* release with repeated release of *t/t* males, resulting in an additional buildup of sterile males. The additional sterile males can push the population beyond the eradication threshold: monthly *t/t* releases of *µ*_t/t_ = 2% and 10% trigger eradication after 25.4 and 6.0 years, respectively (Fig. 1D.2–3). The direct release of *t/t* sterile males is particularly effective at a later stage when each *t/t* male added contributes a larger fraction of a declining population. Fig. 1E explores more systematically the point where *t/t* releases cause the system to switch to population collapse (henceforth *µ*^⋆^): at baseline, the minimal monthly release effort for eradication is *µ*^⋆^ = 1.68%. Eradication time will drop substantially with increasing *t/t* release effort, but will also increase the maximum population size reached during the eradication campaign (Fig. 1E). Repeated *+/t* fertile releases are generally unnecessary, as *t* often reaches equilibrium regardless of initial frequency, and continued release of fertile animals prolongs population persistence.

### Refining releases We explored four strategies to reduce release effort and eradication times (Fig. 2)

#### Trap & release

In this approach, *t* releases are combined with (non-selective) trapping of resident animals (male and female). Trapping strongly reduces required *t* releases and eradication times. For example, trapping 5% residents per month will lower the required annual *t/t* releases by 40% to *µ*^⋆^ = 1.01% and eradication time drops by 31% to 17.5 years at *µ*_t/t_ = 2% (Fig. 2A and Tab. S1). Note that trapping will become less effective as *t* spreads, because *t* carriers born in the population are also removed, thus reducing the difference in *t* frequency between released and trapped animals. Trapping alone will only be sufficient for population eradication without the *t* releases at 40% per month at baseline (Text S2).

**Figure 2.**
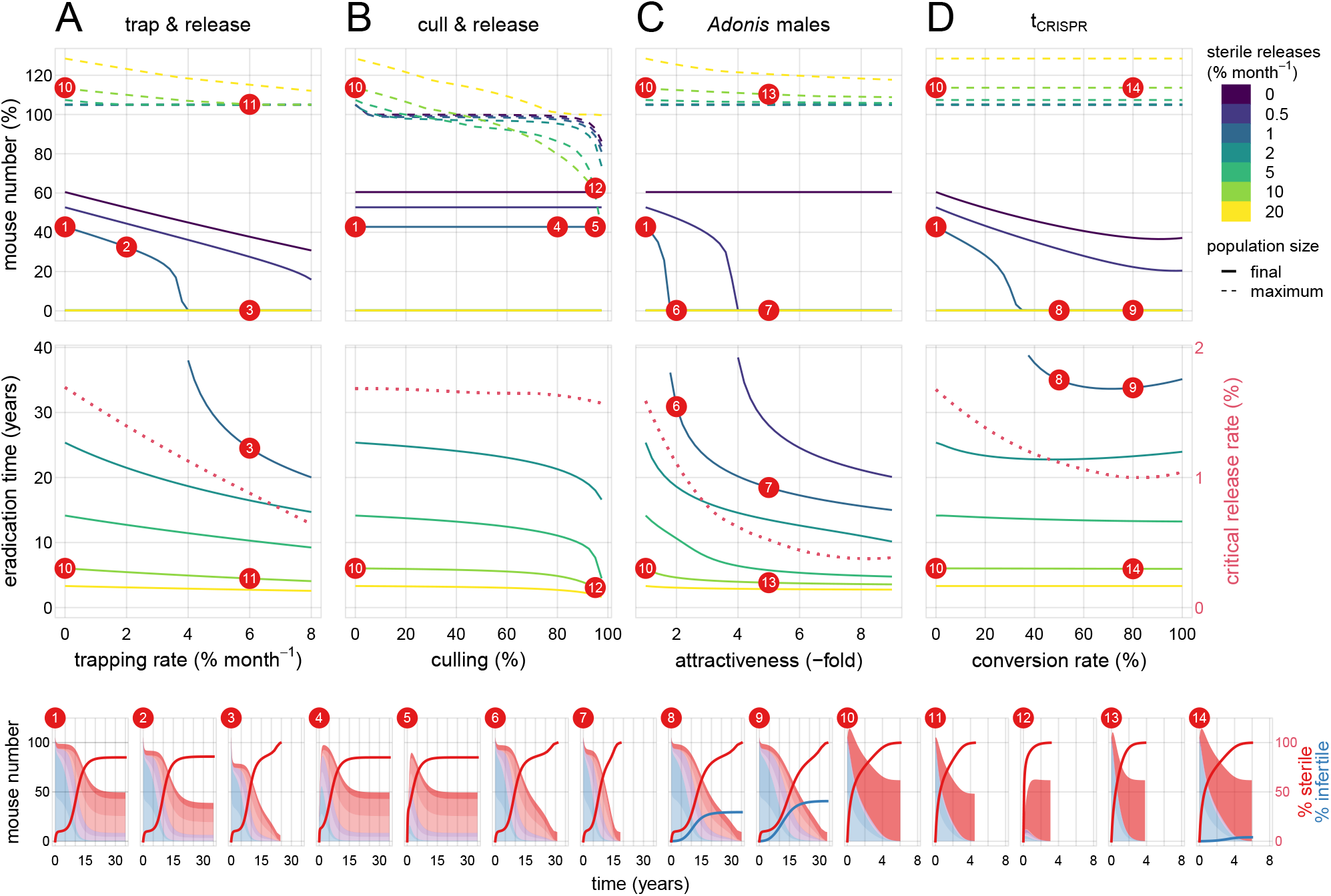
Refining sterile *t* releases. Eradication time and release effort may be reduced using the following refinement strategies: **A**. trap and release, **B**. cull and release, **C**. *Adonis* males, **D**. *t*_CRISPR_. Top row shows maximum (dashed lines) and final (solid lines) target population size, center row eradication time and critical release thresholds. For each method, seven levels of sterile release effort are depicted (colours). Red numbers indicate the scenarios illustrated in the bottom panels (for simplicity, the top row refers to final population sizes for scenarios 1—9 and maximum population sizes for scenarios 10—14). *Bottom row panels:* Population dynamics for 14 scenarios illustrating the effect of each refinement with other parameters held at baseline. Sterile *t/t* release rates are at 1% per month for scenarios 1–9 and at 10% per month for scenarios 10–14. Red and blue lines show the % of sterile males and infertile females, respectively.

#### Cull & release

Here, we examine the scenario where sterile *t* males are released into a population that has previously been culled by other means (for example with rodenticide). In general, the culling has little to no impact on the release effort required for eradication (Fig. 2B & Fig. S4): populations typically recover before the sterile *t* has spread genetically, as ecological dynamics are typically faster than gene frequency changes. That said, culling does reduce eradication times if >90% are culled. For example, with a 95% cull, eradication time will drop from baseline to 18 years (29% reduction). The main appeal of this approach may be that it allows flooding the population with sterile *t/t* males without causing overpopulation: for example, a 95% cull followed by 10% *t/t* releases per month results in complete eradication after 3.1 years, and maximum population size never exceeding 62% of its original size (scenario 12 in Fig. 2).

#### Adonis males

Releasing *t/t* males that are hyper-attractive to females and/or competitive against resident males would be an effective tool to enhance *t* impact. Here, we measure the competitiveness of released *t/t* males with parameter *a* (with *a* = 1 indicating parity and *a* > 1 increased competitiveness) relative to resident males. For example, at threefold competitiveness (a = 3), release rates fall to 0.76% (55% reduction) and eradication time to 16.1 years (37% reduction; Fig. 2C).

#### *t*_CRISPR_

We also explicitly compare our approach to *t*_CRISPR_, currently the leading genetic biocontrol strategy developed for house mice (16). *t*_CRISPR_ is a genetically modified version of the sterile *t*, that, in addition to the effects examined here, carries a CRISPR/Cas cassette disrupting female fertility gene *Prl* during spermatogenesis (16). The CRISPR-mediated conversion, occurring with efficiency *c*, renders homozygous *Prl-/Prl-* females infertile, providing an additional mechanism of reproductive suppression. At 80% conversion efficiency, *t*_CRISPR_ lowers required release effort by 41% to 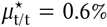 and eradication time by 8% to 23.3 years (Fig. 2D). The reduction in eradication time is smaller than with other refinements, because *t*_CRISPR_ relies solely on genetic spread, which is constrained by generation time. Interestingly, the genetic load imposed by male sterility and female infertility can slow its own spread, thereby limiting population suppression. For example, maximal conversion of 100% does not minimize eradication time (Fig. 2D, also noted by (16)). Similarly, weaker fertility costs in *Prl-* homo- and heterozygotes could, in principle, produce stronger suppression (Fig. S3–4).

### Parameter sensitivity

We assessed model parameter sensitivity of required release effort *µ*^⋆^ and eradication times *t*_erad_ (at *µ*_t/t_ = 2% per month) for five release scenarios (Fig. 3 and Table S1): basic sterile *t* releases, sterile *t* release with 5% trapping, sterile *t* release after a 95% cull, releasing *Adonis* males (with 3-fold competitiveness), and releasing *t*_CRISPR_ (at 80% conversion efficiency).

**Figure 3.**
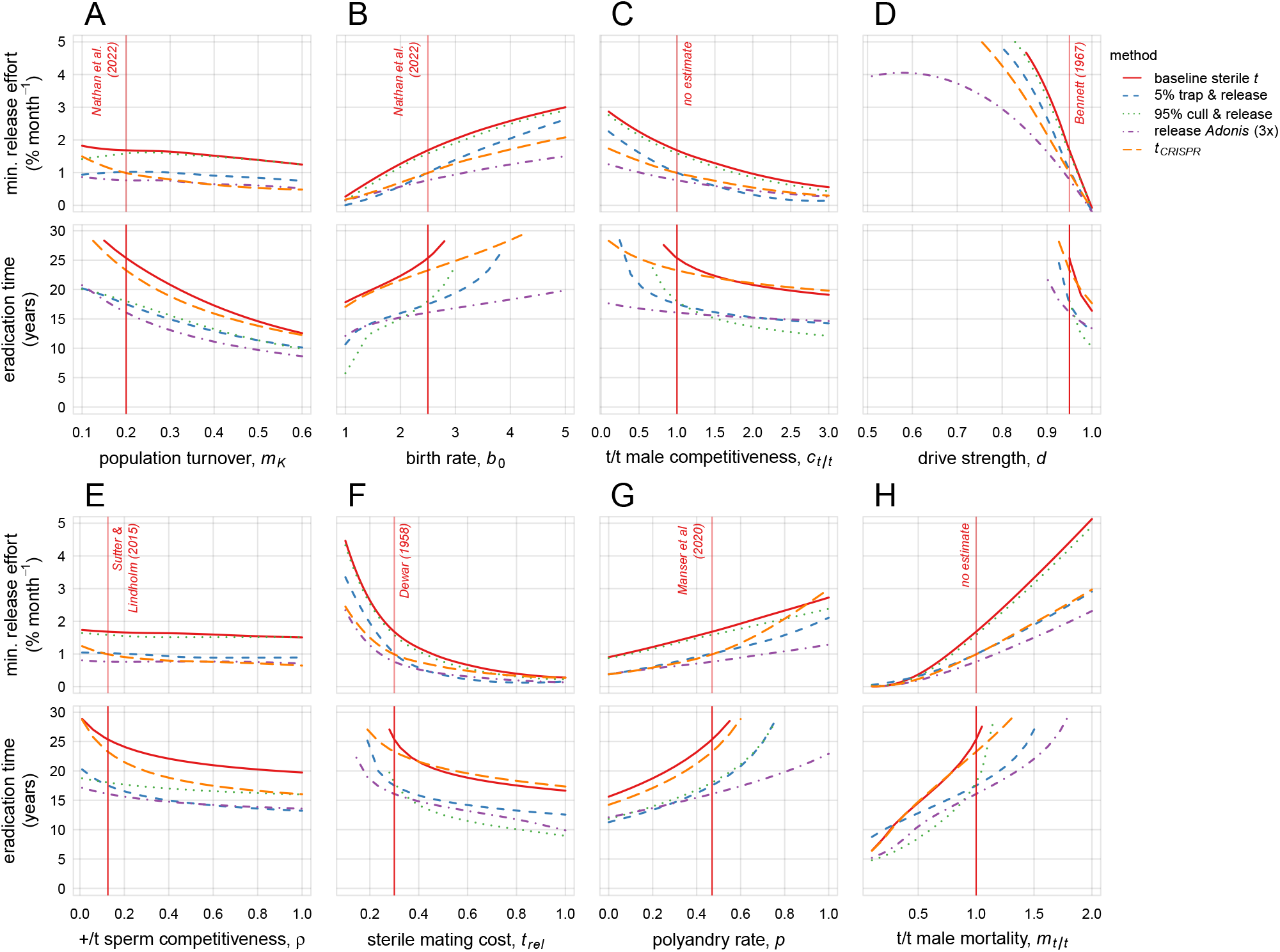
Parameter sensitivity. The impact of eight model parameters on the minimal *t/t* sterile release effort required for eradication (µ^⋆^) and time to eradication (at *µ*_*t*/*t*_ = 2% per month and 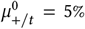). Colors/linetype represent five release strategies: baseline sterile *t* method (red/solid), 5% trap and release (blue/dashed), 95% cull and release (green/dotted), release of *Adonis* males with 3-fold competitiveness (violet/dash-dotted), and *t*_CRISPR_ with 80% conversion efficiency (orange/long-dashed). Minimal release efforts *µ*^⋆^ were smoothed with local regression (LOESS) as they were numerically calculated from a discrete array of release rates. Vertical red lines mark the parameter values used in the baseline scenario (Table 1).

#### Ecology

Population turnover (*m*_*K*_) modestly affects required release effort but strongly accelerates eradication times as lower generation times accelerate genetic spread. A high turnover of 60% per month reduces eradication times to 8.6–12.6 years (42–50% reduction to (method-specific) baselines). Higher birth rates (*b*_0_) increase release effort and eradication times. For example, doubling birth rates from 2.5 to 5 pups per month increases *µ*^⋆^ by 79– 160%, and eradication times become prohibitively large for *µ*_t/t_ = 2%.

#### Male reproduction

Gene drive strength is pivotal. Values below 85% generally preclude eradication for almost all methods, whereas near-complete drive of 100% reduces release effort to almost zero. *t/t* male competitiveness (born and released, as opposed to *Adonis* males attractiveness which *only* affects released *t/t* males) critically affects *µ*^⋆^ and *t*_erad_. Surprisingly, *+/t* sperm competitiveness has little effect, particularly on *µ*^⋆^, possibly due to frequency-dependence (see Fig. S2).

#### Female reproduction

Sterile mating costs strongly influence outcomes: low costs ( 10%) increase release needs by 149–253%, whereas high costs (>50%) drastically reduce release requirements to 0.1-0.3% per month (76–83% reduction). Polyandry increases *µ*^⋆^ (51-109% increase at 100% polyandry) and eradication times, with *t*_CRISPR_ particularly sensitive due to its reliance on genetic spread (209% increase at *p* = 100%).

#### Survival

Viability selection against *t/t* males significantly impacts both *µ*^⋆^ and *t*_erad_; survival costs >50% make eradication unfeasible in terms of eradication times, whereas very high *t/t* survival reduces *µ*^⋆^ to nearly 0 and eradication times at *µ*_t/t_ = 2% to under 10 years.

### Shiny app

To enable both reproduction of the scenarios presented here and exploration of parameter settings beyond, we provide an interactive R Shiny application that allows users to adjust model parameters and compute outcomes in real time (see https://shiny.math.uzh.ch/user/manser/sterilet/).

## Discussion

The aim of our model was to assess feasibility of using the naturally occurring, sterility-inducing *t* haplotype to control invasive house mouse populations. Our model suggests that a combination of early *+/t* fertile male releases, initiating genetic *t* spread, and repeated *t/t* sterile male releases, raising sterility directly, can principally cause population collapse. However, the strategy typically requires years of continued releases to gradually elevate the frequency of sterile males to levels sufficient for population control. Its effectiveness could be considerably enhanced if *t* releases are combined with trapping, culling or the release of highly competitive males, reducing release numbers, population burden and eradication times. Importantly, the approach critically hinges on key aspects of mouse reproductive biology, demography, and behaviour, many of which remain poorly understood. Targeted empirical research will be essential to robustly assess its potential as a control tool.

A central open question concerns the reproductive costs, and thus demographic impact, imposed by sterile males, as even few fertile males may sustain population persistence. In our model, we represented sterile male costs as delays in the female reproductive schedule. Laboratory evidence shows that mating with vasectomised males induces pseudopregnancy, delaying return to estrous for 10–19 days (40; 38; 41; 42). Whether this occurs in wild house mice, whether *t/t* sterile males trigger the same hormonal response, and how such delays compare to baseline interbirth intervals remains to be investigated.

A key limitation of our model is that it assumes a panmictic population for computational tractability, allowing us to explore feasibility across a broad range of parameters. Spatial structure is expected to play a central role in the effectiveness of releases. On one hand, small and disconnected demes will be more susceptible to stochastic loss of fertile males, thus causing local population extinction. Note that deterministic *t* frequency equilibria of ~90% are relatively close to this *t* fixation (i.e. extinction) boundary (43). Accordingly, previous modelling work on the system reports elevated extinction risk under high population structure (43; 44). On the other hand, the reproductive interference by sterile males will be spatially local rather than global (as assumed here), potentially leaving reservoirs of fertile males that could recolonise locally depleted patches, thereby slowing overall suppression dynamics. Birand et al. (45; 16) have developed a spatially explicit modelling framework, and found that eradication probability increased slightly with larger dispersal distances for both sterile *t* and *t*_CRISPR_ (16). Overall, the dynamics of their spatially explicit models are qualitatively consistent with those of our panmictic model. Incorporating spatial and temporal complexities of given target populations would be an important next step should the quantification of key parameters identified in this study suggest the general feasibility of the approach.

It is further unknown how females will respond to sterile *t/t* males, both when fertile alternatives are available and when all socially and spatially available males are sterile. There is also conflicting evidence whether females avoid *t* carrying males. While female *t* avoidance would be adaptive, empirical results from lethal *t* haplotypes are mixed: some studies report olfactory avoidance of *+/t* males (46), whereas others find no evidence for discrimination (30; 47). A related question is how social dynamics will shift under extreme conditions near the end of a campaign, when populations become small and heavily male-biased. Elevated male–male aggression could undermine the effectiveness of releases. In our model, we assumed a (slight) decrease in mortality with decreasing population density, but did not account for possible effects of extreme sex ratios and low density on other parameters. For example, both density and sex-ratio are likely to influence polyandry: while remating rates tend to decline at low density (27), an excess of males may intensify sperm selection against *+/t* males.

Taken together, the uncertainties regarding mouse biology, ecology, and behaviour under relevant field conditions limit an accurate assessment of the sterile *t* and related methods. Moreover, much existing evidence comes from European or North American populations where mice live commensally with humans, which may differ substantially from feral island populations where mice occupy less predictable, non-anthropogenic habitats (48; 49).

A practical limitation of the sterile *t* method is the need for large and repeated male releases (1.7% of carrying capacity per month at baseline). The success of sterile male techniques in insects stems from the ability to flood populations with sterile males at scale (23; 50). While the continued release of *t/t* sterile males is very effective to reduce eradication time (in extreme scenarios to under <5 years, Fig. 1E), it is unclear whether this is feasible in a mammal species such as the house mouse. The challenge is not breeding capacity: *t*-carrying mice are readily available in laboratory stocks and could be bred cheaply and in large numbers (e.g. from crosses or enclosures of *+/t* males and *t/t* females). The bottleneck is whether released animals are able to settle into an population with an established social structure. Genetic studies of island populations show remarkably small signatures of introgression despite continual introductions, suggesting that settlement may be difficult (51). Despite this, an early attempt to release *t*-carrying mice into an island population reported that the *t* haplotype persisted for several years after release (52). Timing and spatial distribution of releases may boost settlement and overall effectiveness (53). Perhaps more importantly, the transient burden caused by continued releases may be ecologically unacceptable. Combining releases with extensive trapping, or following population reduction by other means, could both facilitate settlement and avoid transient increases in population size resulting from large-scale releases, but would entail substantial additional material and labour costs. Trapping may be necessary regardless to monitor eradication progress (but see Piaggo et al. (54) for a method that does not require animal capture). Culling does not substantially reduce the release threshold required for eradication, but could enable largescale *t/t* releases without imposing an additional population burden. More broadly, integrating genetic biocontrol with conventional population management may improve the feasibility of eradication, particularly during the critical “last mile” when populations have already been substantially reduced (55).

The *Adonis* strategy illustrates the potential to improve efficacy without additional field labour, although its effectiveness remains speculative, as the traits underlying male attractiveness and competitiveness remain poorly understood. Simply releasing males with larger body size, a trait that varies drastically in natural populations, may boost male competitiveness. In insects, aromatherapy is used to increase mating performance of sterile males up to threefold (56). Volatile pheromones such as ESP1 have been shown to upregulate female sexual behaviour (57). Note that the *Adonis* effect could be even greater if both *t/t* sterile and *+/t* fertile males outperform resident males (we modelled only *t/t* males). However, if competitiveness is heritable and eradication fails, this could produce a more robust pest population. To maximise settlement, mating success, and survival, it would generally be advisable to backcross sterile *t* mice onto the local genetic background before release, thereby minimising genotypic and phenotypic differences with resident mice, potentially even conferring a hybrid vigour advantage (58). Finally, combining refinement strategies could improve efficacy in a non-additive manner. For instance, trapping may accel-erate population suppression using *t*_CRISPR_, which heavily relies on rapid turnover.

An important consideration is how the sterile *t* approach compares with engineered alternatives such as *t*_CRISPR_. In our explicit model comparison, sterility from the naturally occurring *t* already accounts for much of the population suppression achieved by *t*_CRISPR_, with the engineered addition of female infertility primarily reducing the number of releases required (−41% at baseline), but only marginally impacting eradication time (−8% at baseline). Several of the refinements considered here may similarly reduce release requirements without modifying the naturally occurring *t* haplotype. Other engineered gene-drive strategies may achieve more rapid suppression: modelling of homing and X-shredding drives, for example, predicted more rapid eradication of island mouse populations (45; 16). However, despite experimental efforts, developing functional constructs of these types appears challenging in house mice (59). Thus, the relative merits of natural and engineered strategies may depend not only on their population-suppression potential, but also on the feasibility, cost, and complexity of developing and deploying them at scale.

If feasible, the sterile *t* method may offer several advantages over current control methods and synthetic gene drives. Population suppression occurs through reduced reproduction rather than killing resident mice using toxicants, which has raised animal welfare concerns. The *t* haplotype is restricted to *Mus musculus*, making the method species-specific, thus limiting direct effects on non-target species. This could make the approach relevant in human-inhabited areas where toxicant-based control is problematic (8). The sterile *t* method harnesses a naturally occurring genetic element already present in laboratory and wild populations, potentially enabling relatively simple and low-cost breeding and release. Importantly, *t* haplotypes have maintained a strong drive level despite millions of years of coevolution and genomic conflict with their host, suggesting that the evolution of drive resistance may be less likely than in synthetic systems, where resistance has been reported (60). Finally, the method does not require genetic modification, which may reduce regulatory and ethical barriers to its testing and use.

## Material and Methods

### General approach

To assess the sterile *t* method as a pest control tool, we model release programmes of *t* carrying males into a target population. We track genotype abundances at time *t* with the vectors

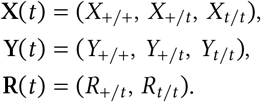

for females, born males, and released males, respectively (note that boldface denotes vectors). Relative genotype frequencies are then

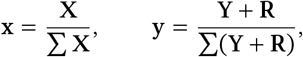

where, for convenience, we gather released and born males in a single vector **Y** + **R** = (*Y*_+/+_, *Y*_+/*t*_ + *R*_+/*t*_, *Y*_*t*/*t*_ + *R*_*t*/*t*_). The overall population size at time *t* is simply *N* = ∑ **X**+∑ **Y**+∑ **R**. Changes in the absolute and relative abundance of the genotypes, and ultimately the ability of released *t* animals to survive and breed in the target population, will depend on the biological processes described below. Overall, we model the dynamics of the system in the following set of eight coupled differential equations.

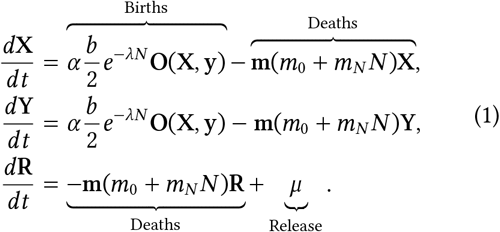

Here, 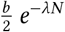 and (*m*_0_ + *m*_*N*_*N*) define the demographic profile (i.e. density dependent birth and death rates) in the target population (see section A). Vector *µ* describes the release of *t* carrying males (*+/t* and *t/t*) into the target population (see section B). Parameter α critically measures the *impact* of *t/t* sterile males on offspring production (see section C). Vector **O**(**X, y**) counts the (absolute) number of the three offspring genotypes produced, which will depend on the female abundances **X**, relative male frequencies **y** as well as a number of pre- and postmating selective processes: multiple mating rate *p*, the competitiveness of male genotypes **a**, sperm competitiveness of *+/t* males *ρ*, as well as drive strength *d* (see sections D and E). Finally, vector **m** measures the differential mortality rates of the three genotypes (see section F).

### A. Ecology

*Birth and death*. Ecological assumptions are illustrated in Fig. 1B. We start the model with a target population of wildtype animals *+/+* at carrying capacity *K* and an even sex ratio. Females give birth to offspring at baseline birth rate *b*_0_ per month, of which a fraction *m*_pup_ die during the time until weaning due to pup mortality, giving us an recruitment rate of *b* = *b*_0_(1 − *m*_pup_). Recruitment rate decreases exponentially (to avoid negative recruitment) with population density due to increased competition with density-dependent decay parameter λ. At population density *N*, the overall recruitment rate is thus given as *b*(*N*) = *b e*^−λ*N*^. Equivalently, we assume that all animals die with a baseline mortality rate of *m*_0_ per month, which increases linearly with population density defined by density-dependent mortality rate *m*_*N*_, such that mortality rate at density *N* is given as *m*(*N*) = *m*_0_ +*m*_*N*_ *N*.

*Carrying capacity*. The population reaches carrying capacity *K* when number of females (or males) produced equals the number that die: 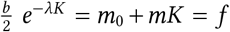. Here, *f* denotes the turnover rate (fraction of animals born/dead per month) at carrying capacity. The factor 1/2 accounts for the fact that only half of the offspring are female (or male). For example, if *f* = 0.4, 40% of animals are ‘replaced’ each month when the population is at carrying capacity.

Because these quantities are more biologically meaningful, we will henceforth describe the demographic profile of the target population via its carrying capacity *K*, baseline birth *b*_0_, adult and pup mortality (*m*_pup_ and *m*_0_), and turnover at carrying capacity *f*. The parameters *λ* and *m*_*N*_ defining density-dependence are chosen such that births and deaths intersect at *K* : 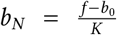 and 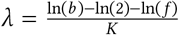.

### B. Releases

Fertile *+/t* and sterile *+/t* drive males are released into the population either once or continuously. Parameters 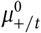 and 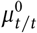 define the number of animals released at the beginning of a release campaign relative to carrying capacity *K*. Release vector *µ* = (µ_+/*t*_, *µ*_*t*/*t*_) measures the number of *+/t* and *t/t* males, respectively, released continuously per month. For example, if *µ*_*t*/*t*_ = 0.05 and *K* = 1000, 50 *t/t* males are released into the target population every month.

### C. Sterility costs

The central tenet of the sterile *t* method is for *t/t* sterile males to hamper reproductive output *b* by raising the probability of unproductive matings *P*_0_, which occur when females *exclusively* mate with *t/t* sterile males during a given estrous. It is clear that female reproductive rates will decline from normal levels to zero as the frequency of sterile males increases from 0 to 1, thus the probability of unproductive matings will go from *P*_0_ = 0 to *P*_0_ = 1. The precise impact of sterile males on reproduction, particularly for intermediate frequencies (0 < *y*_*t*/*t*_ < 1), however, is so far unknown and will crucially depend on the (reproductive) costs imposed by unproductive matings. The impact of sterile males on reproduction will be a composite measure of the probability of an unproductive mating and the reproductive repercussions (cost) incurred by such an event.

We here conceptualise the (reproductive) cost incurred by unproductive matings in terms of the relative time delay they cause, defined by parameter *t*_*miss*_. Specifically, if *t*_*IB*_ is the average time between two births (interbirth interval) for ‘typical’ females in the absence of sterile males, we define parameter *t*_*miss*_ as relative time added *every time* a female has an unproductive mating. *t*_*miss*_ can thus be thought of as a time penalty for not finding fertile males, causing a delay in reproduction. Minimally, the time penalty incurred will be 4 days, the duration of an estrous cycle. Studies indicate however that mating with sterile males reliably induces a state of pseudopregnancy in about 80% of females (61), which will increase the time until a female will return to estrous by up to 12–19 days (38; 31). The impact of sterile males on the relative reproductive output α will then be the product of the probability of an unproductive mating *P*_0_ and the time penalty incurred each time a female was unsuccessful, *t*_*miss*_. The relative reproductive output α is then given by the expected share of time spent producing offspring over the total time:

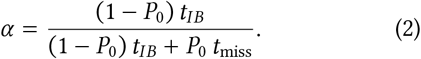

Equivalently, we can define sterile costs as the delay relative to a normal interbirth-interval *t*_rel_ = *t*_miss_/*t*_*IB*_, we have 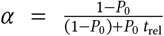. Figure 1C illustrates how α decreases with *P*_0_ for different *t*_rel_. With small *t*_rel_, repro-ductive rates remain close to baseline until sterile males become very frequent. By contrast, larger *t*_rel_ values cause more steady declines. In the special case *t*_rel_ = 1 (*t*_miss_ = *t*_*IB*_), females miss an entire reproductive cycle in case of an unsuccessful mating, so α decreases linearly with *P*_0_. Note that the probability of an unproductive mating *P*_0_ will generally increase with the frequency of sterile males in the population *y*_*t*/*t*_. However, due to additional processes such as male-male competition, female preference and polyandry, the relationship may not be linear (see next section).

### D. Mating

In this step, now that we have defined the reproductive repercussions of unproductive matings, we need to identify the factors that will determine the probabilities that a female will mate with the various male genotypes, and with it the probability of unproductive matings *P*_0_.

#### Premating selection

First, males compete for mating opportunities. Let vector **a** = (a_+/+_, a_+/*t*_, a_*t*/*t*_) measure the competitiveness of the three male genotypes in premating sexual selection, which we think of as a composite measure of their ability to compete against rival males as well as to attract females as mating partners (female preference). In particular, *c*_+/*t*_ and *c*_*t*/*t*_ measure the competitiveness of *+/t* and *t/t* males relative to resident wildtype *+/+* males, whose competitiveness equals unity.

#### Polyandry

We assume that females either mate once (at probability 1 − *p*) or twice (with probability *p*) during an estrous cycle. Polyandry is an important process here for two reasons. First and crucially, it will impact on the probability of unproductive matings *P*_0_, as the likelihood of encountering at least one fertile male will increase with every additional mating partner. Second, it has been repeatedly demonstrated that, as a direct consequence of the sperm manipulation mechanism that causes the gene drive effect, *+/t* heterozygote males are severely compromised in their sperm competitive ability against rival *+/+* males (35; 36; 37). Polyandry will expose *+/t* males to sperm competition against rival males, jeopardising their fertilisation success and thus the genetic spread of *t* via *+/t* males (see below for a description of the fertilisation process).

#### Mating

We now have all the necessary information to calculate the frequency of the 9 ordered monandrous combinations 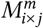 as well as the 27 ordered polyandrous mat-ing combinations 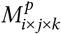, between females of genotype *i* and males of genotype *j* and (in the case of polyandry) *k*. At this point, we also add the released males **r** = (*r*_+/*t*_, *r*_*t*/*t*_) to the male mating pool. We have

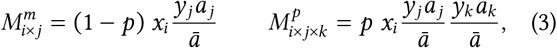

where ā denotes the mean competitiveness across all male genotypes. Moreover, we can now also calculate the probability of unproductive matings, which occurs whenever a female *only* encounters *t/t* males.

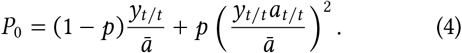

### E. Offspring

Now that we know the probabilities of the mating combinations, we can calculate the proportions of zygote genotypes they produce. Let *l* ∈ {+/+, +/*t, t*/*t*} denote the offspring genotypes. Each monandrous mating combination *i*×*j* produces offspring proportions 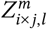, while polyandrous combinations *i* × *j* × *k* produce 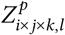. Zygote production is governed by three biological processes:

#### Male sterility

Mating combinations that *only* involve *t/t* sterile males will not produce any offspring. We further assume that there are no fertility differences between *+/+* and *+/t* males, and that polyandrous females restore fertility completely if they mate with at least one fertile male (*+/+* or *+/t*).

#### Gene drive

We characterise the strength of gene drive with parameter *d*, which measures the proportion of *t* that fertile *+/t* males transmit to their progeny. Hence, *d* = 0.5 corresponds to Mendelian segregation, while *d* = 1 represents complete drive, where *+/t* transmit the *t* to all offspring. Parameter *d* can be thought of as measure of fitness of *t* sperm (relative to *+*) *within* a *+/t* male ejaculate (intragenomic competition).

#### Sperm competition

Parameter *ρ* measures the sperm competitive ability of *+/t* males relative to resident *+/+* males, whose sperm competitive ability equals unity. If *ρ* = 1, *+/t* and *+/+* males do not differ in their sperm competitive ability, such that both males would secure half of the fertilisations in a polyandrous mating. If *ρ* < 1, *+/t* have reduced sperm competitiveness, such that they only fertilise 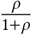 eggs when in sperm competition against *+/+* resident males. Parameter *ρ* can be thought of as a measure of fitness of a *+/t* male’s sperm (relative to *+/+* males) in sperm competition *between* males.

#### Offspring genotypes

Knowing the frequencies of each mating combination **M** and the proportions of zygotes they produce, **Z**, we can calculate the number of offspring of each genotype at any given time as the vector **O** = (*O*_+/+_, *O*_+/*t*_, *O*_*t*/*t*_). In matrix notation, offspring genotypes are obtained by multiplying the mating vector with the corresponding zygote matrix **O** = **M**^*m*^**Z**^*m*^ + **M**^*p*^**Z**^*p*^, where each row of **Z**^*m*^ and **Z**^*p*^ corresponds to a mating combination, and each column corresponds to an offspring genotype *l*.

### F. Survival

Our model allows mortality rates to be sex- and genotype-specific. We denote the relative mortality rates by 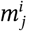, where *i* ∈ {*y, x*} indicates sex and *j* ∈ {+/+, +/*t, t*/*t*} indicates genotype. Wildtype animals define the baseline, such that 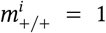. For example, 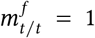 indicates that *t/t* females have the same mortality as wildtype females, while 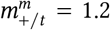 corresponds to a scenario where *+/t* males die at a rate 20% higher than their *+/+* counterparts. The absolute mortality at population size *N* is then given by the vector **m**^*i*^ (*m*_0_ + *m*_*N*_ *N*).

Note that this formulation assumes that relative mortality differences are independent of population density.

### Refining release campaigns

We modelled trapping by adding terms **−τX** and **−τY** to our differential equations 1), such that males and females born in the population (irrespective of their genotype) are removed at a constant rate τ. With cull and release, we simply start the population at a fraction γ of carrying capacity, such that *N*_0_ = γ *K*. The *Adonis* strategy was implemented by introducing an attractiveness factor a for released (only) sterile *t/t* males, analogous to equation (3). To model *t*_CRISPR_, we extended the system by adding an additional locus with two alleles (*Prl* and *Prl-*), yielding nine differential equations per sex instead of three (see R code).

### Parameter sensitivity

Critical release rates were determined numerically by calculating dynamics across a grid of release rates (µ_*t*/*t*_ = [0, 0.4] with grid resolution 0.01; 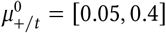 with grid resolution 0.05), identifying the scenario requiring the fewest released males as *µ*^⋆^. We considered a population eradicated if number of fertile females fell below 0.1% of carrying capacity *K*. Model parameters were varied one at a time.

### Implementation

The full system of differential equations is beyond analytical tractability. The results presented here are generally based on numerical solutions using the ode package in R.

## Acknowledgements

We thank Bruce Hay for very insightful feedback on a previous version of the manuscript. We thank our funding bodies for the support: Swiss National Science Foundation (SNF) Spark grant CRSK-3_190749 (A.M.) and project grant 310030_207607 (A.M. and A.K.L.).

