## supplementary material for "Assessing a natural gene drive for population control in house mice"

#### Texts

**S1 Calculating the frequency of sterile males required for eradication** 2

**S2 Calculating the trapping required for eradication without *t* releases** 2

#### Figures

S1 Calculating the frequency of sterile males required for eradication . . . 4
S2 Sterile *t* frequency equilibria for model parameters without release . . . 5
S3 Focus: Polyandry . . . 6
S4 Focus: Cull & release . . . 7
S5 Focus $t_{CRISPR}$: Parameter dependence . . . 8
S6 Focus $t_{CRISPR}$: Maximising popoulation suppression . . . 9

#### Tables

S1 Parameter sensitivity . . . 10

### S1 Calculating the frequency of sterile males required for eradication

The full system of differential equations is (likely) not analytically tractable. We can however calculate the frequency of sterile male required for population decline, denoted as $y^\star_{t/t}$. The population will decline whenever the total number of females $X = \sum \mathbf{X}$ (eq. 1) satisfies

$$\frac{dX}{dt} = \alpha \frac{b}{2} e^{-\lambda N} - (m_0 + m_N N)X < 0. \tag{S1}$$

To obtain an analytical expression, we treat the population size $N$ as constant and therefore do not need to track the number of released males $R$ or newborn males $Y$ explicitly. Substituting $\alpha = \frac{1-P_0}{(1-P_0)+P_0\, t_{\mathrm{rel}}}$ (eq. 2) and $P_0 = (1-p)\frac{y_{t/t}}{\bar{a}} + p\left(\frac{y_{t/t} a_{t/t}}{\bar{a}}\right)^2$ (eq. 3) into the condition $\frac{dX}{dt} < 0$ yields a quadratic equation in $y_{\mathrm{t/t}}$. This gives us the critical sterile male frequency

$$y^\star_{t/t}(N) = \frac{bp - b + A(p-1) \pm \sqrt{(b+A)(b(1+p)^2 + 2e^{\lambda N}(m_0 + m_N N)(t_{\mathrm{rel}}(p-1)^2 - (p+1)^2))}}{2p(b+A)}, \tag{S2}$$

where, for compactness, we define $A = 2e^{\lambda N}(m_0 + m_N N)(t_{\mathrm{rel}} - 1)$.

The first-order derivative $\frac{dy^\star_{t/t}(N)}{dN}$ shows that $y^\star_{t/t}(N)$ monotonically decreases with respect to $N$ across biologically relevant parameter values ($\lambda, m_N, m_0, t_{\mathrm{rel}} > 0$). We can therefore obtain a simplified upper bound for the critical frequency by evaluating the expression at $N = 0$. We have

$$y^\star_{t/t} = \frac{p-1}{2p} + \frac{\sqrt{(b + 2m_0(t_{\mathrm{rel}} - 1))((b - 2m_0)(1+p)^2 + 2m_0(p-1)^2 t_{\mathrm{rel}})}}{2p(b + 2m_0(t_{\mathrm{rel}} - 1))}. \tag{S3}$$

For the baseline scenario (Table 1), we have $y^\star_{t/t} = 95.2\%$, thus $t/t$ sterile need to exceed this proportion to cause population declines at $N = 0$.

We can put the required sterile frequency $y^\star_{t/t}$ in relation to the equilibrium frequency of steriles $\hat{y}_{t/t}$. Dunn and Bennett (1) calculated in the absence of polyandry and viability selection as

$$\hat{y}_{t/t} = 4(d - \frac{1}{2})^2 \tag{S4}$$

Population eradication without repeated releases is expected if $\hat{y}_{t/t} > y^\star_{t/t}$. Substituting eq. S4 into this inequality and solving $d$, we get

$$d^\star > \frac{1}{2} + \sqrt{\frac{y^\star}{4}},$$

which means that, under baseline scenario and monandry, the target population will be eradicated after a single release if $d^\star > 98.8\%$.

### S2 Calculating the trapping required for eradication without *t* releases

We can calculate the critical trapping rate $\tau^\star$ that will result in population collapse in the absence of sterile $t$ releases. In the absence of releases, we have

$$\frac{dX}{dt} = \frac{b}{2} e^{-\lambda N} X - (m_0 + m_K N)X - \tau X$$

with $X = \sum \mathbf{X}$ and $N = 2X$. To calculate the stability of the extinction equilibrium $\frac{dX^\star(t)}{dt} = 0$ by assessing the sign of the first-order derivative at $X^\star = 0$:

$$\frac{dX'(t)}{dt} = X(\frac{b}{2} e^{-2\lambda X} - m_0 - \tau) - 2m_K X^2.$$

Extinction thus become stable if

$$\tau^\star > \frac{b}{2} - m_0, \tag{S5}$$

which corresponds to a trapping rate of 40% per month in our baseline scenario.

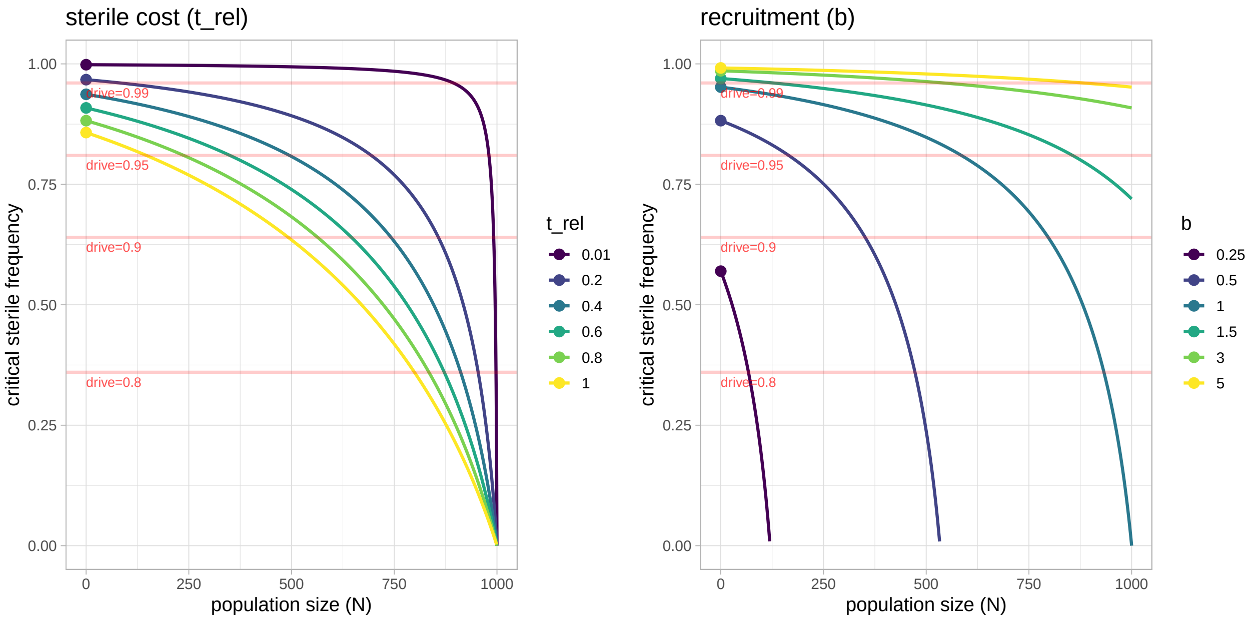


**Figure S1. Frequency of sterile males required for eradication.** Critical sterile frequency $y^{\star}_{\mathrm{t/t}}$ necessary for population suppression (eq. S2). $y^{\star}_{\mathrm{t/t}}(N)$ is shown as a function of population size $N$ and for varying levels of sterile cost $t_{\mathrm{rel}}$ (left) and recruitment $b$ (right). Dots correspond to $y^{\star}_{\mathrm{t/t}}$ from eq. S3. Red horizontal lines show the expected equilibrium *t/t* sterile frequency $\hat{y}_{\mathrm{t/t}}$ according to Dunn et al. (1) in the absence of continued release. The population will be eradicated in the absence of repeated *t* releases if $\hat{y}_{\mathrm{t/t}} > y^{\star}_{\mathrm{t/t}}$. For example, for $d = 0.95$, sterile frequencies will not be high enough for population extinction without repeated, direct release of *t/t* sterile males, but would cause population extinction for $b = 0.25$.

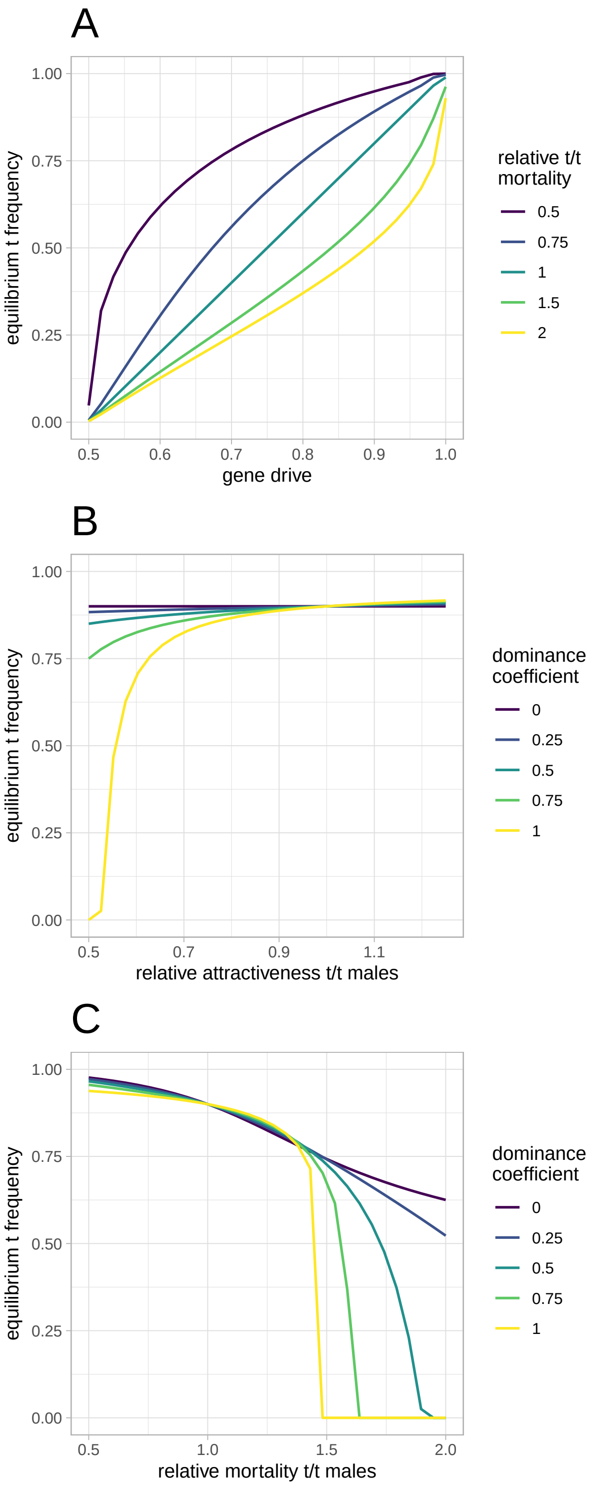


**Figure S2. Sterile *t* frequency equilibria for model parameters without release.**
**A.** Effect of gene drive strength ($d$) and relative *t/t* male mortality ($m_{t/t}$). Equilibrium *t* frequency increases from 0% to 100% as gene drive increases from 50% (Mendelian inheritance) to 100% (complete drive). Mortality determines the shape of this relationship: linear for $m_{t/t} = 1$ (1), concave for $m_{t/t} < 1$, and convex for $m_{t/t} > 1$.
**B.** Effect of male attractiveness. Here, $a_{t/t}$ denotes the attractiveness of sterile *t/t* males relative to *+/+* males (with $a_{+/+} = 1$), while the dominance coefficient $h_a$ determines the attractiveness of *+/t* males as $a_{+/t} = 1 - h_a(1 - a_{t/t})$. As expected, the attractiveness of sterile *t/t* males does not affect equilibrium *t* frequencies (as they do not reproduce), but they do have ecological impact (main Fig. 3). In contrast, the attractiveness of fertile *+/t* males does affect *t* spread, with the *t* disappearing if +/t male attractiveness is roughly halved ($h_a = 1$, $a_{t/t} = 1$), compensating for the (near) two-fold advantage of drive.
**C.** Effect of male mortality. Here again, $m_{t/t}$ denotes the mortality of sterile *t/t* males relative to *+/+* males (with $m_{+/+} = 1$), while the dominance coefficient $h_m$ determines the mortality of *+/t* males as $m_{+/t} = 1 - h_m(1 - m_{t/t})$. Mortality of sterile *t/t* males alone ($h_m = 0$) has little effect on equilibrium outcomes: although the proportion of *t/t* males decreases slightly when they are outlived by other genotypes, steriles do by definition not influence *t* spread. In contrast, mortality costs in fertile *+/t* males strongly affect *t* dynamics; for example, *t* extinction is predicted when *+/t* males experience a 50% survival cost ($m_{t/t} = 1.5$, $h_m = 1$), again compensating the two-fold drive advantage. Remaining parameters: drive $d = 0.95$, polyandry $p = 0$ (Fig. S2).

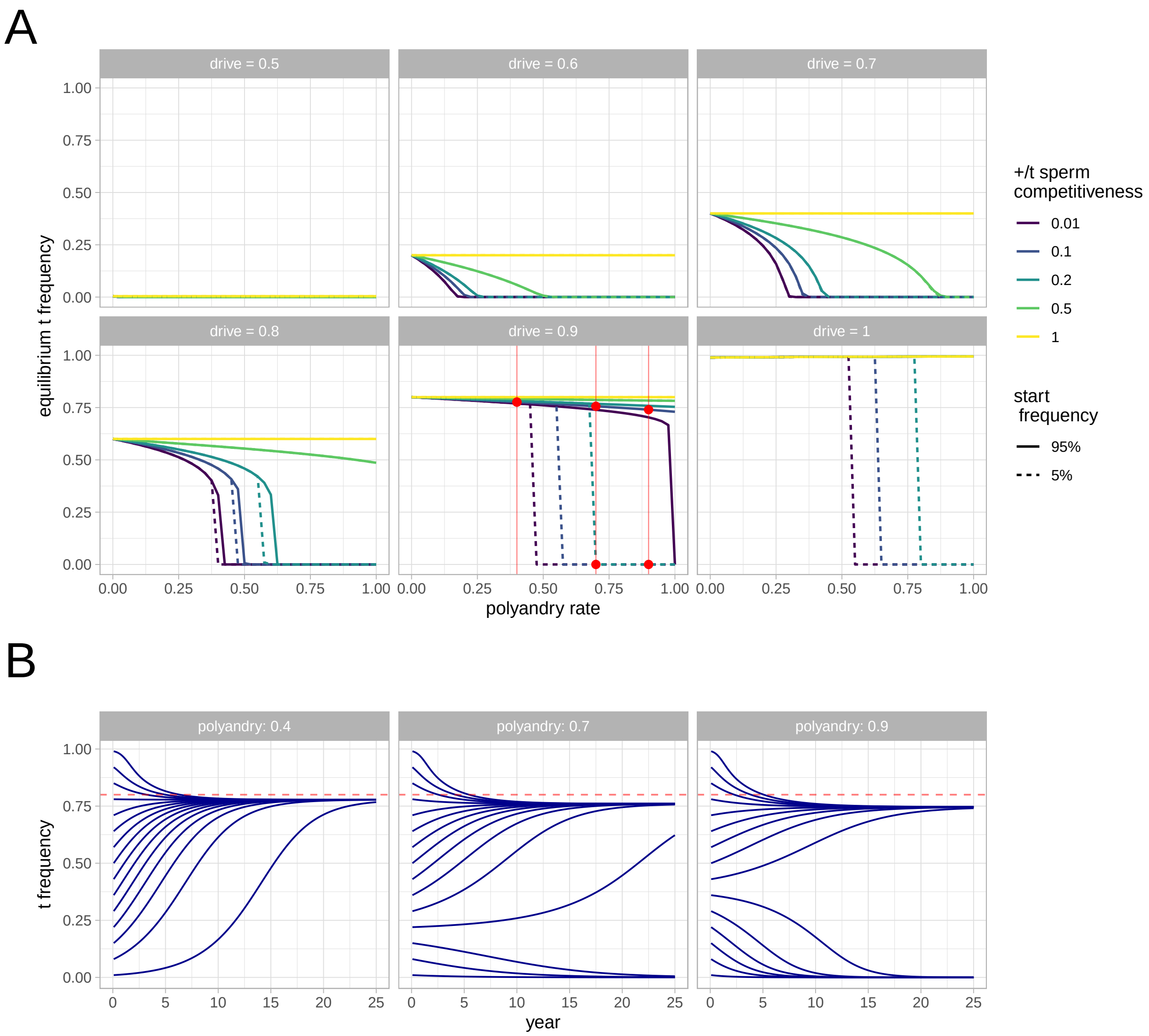


**Figure S3.** **Polyandry: Negative frequency-dependent selection causes invasion thresholds**
**A.** Equilibrium $t$ frequency as a function of polyandry frequency $p$ (x axis), $+/t$ male sperm competitiveness $\rho$ (colors), gene drive $d$ (panels), and $t$ starting frequency (solid: 95%, dashed: 5%). At low drive $d$, polyandry suppresses $t$ frequency in a gradual manner, analogous to the lethal $t$ haplotype version (2). At high drive $d$, however, $t$ frequency remains close to the monandry equilibrium, followed by a sudden phase transition to $t$ extinction. In this regime, equilibrium outcomes become dependent on the starting frequency: low initial $t$ frequencies lead to $t$ extinction, whereas high initial frequencies produce a stable polymorphic equilibrium. Sudden phase transitions and invasion thresholds arise from the frequency-dependent nature of sperm selection, as illustrated in panel B. Red dots and vertical lines correspond to the parameter combinations shown in panel B.
**B.** $t$ frequency trajectories for three levels of polyandry ($p = [0.4, 0.7, 0.9]$) and $+/t$ male sperm competitiveness at baseline ($\rho = 10\%$). For medium to high polyandry frequency, $t$ genetic spread is dependent on starting frequency: below a critical frequency threshold, where $+/t$ males are predominantly exposed to superior $+/+$ males, the $t$ can not spread. Above the threshold, $+/t$ compete more frequently against other $+/t$ or $t/t$ males, where their low competitiveness does not matter. As a result, $t$ spreads to a polymorphic equilibrium. The red dashed horizontal line shows the analytical prediction by Dunn (1) of $\hat{p}_t = 2d - 1$ based on gene drive and $t/t$ male sterility (at drive level $d = 90\%$).

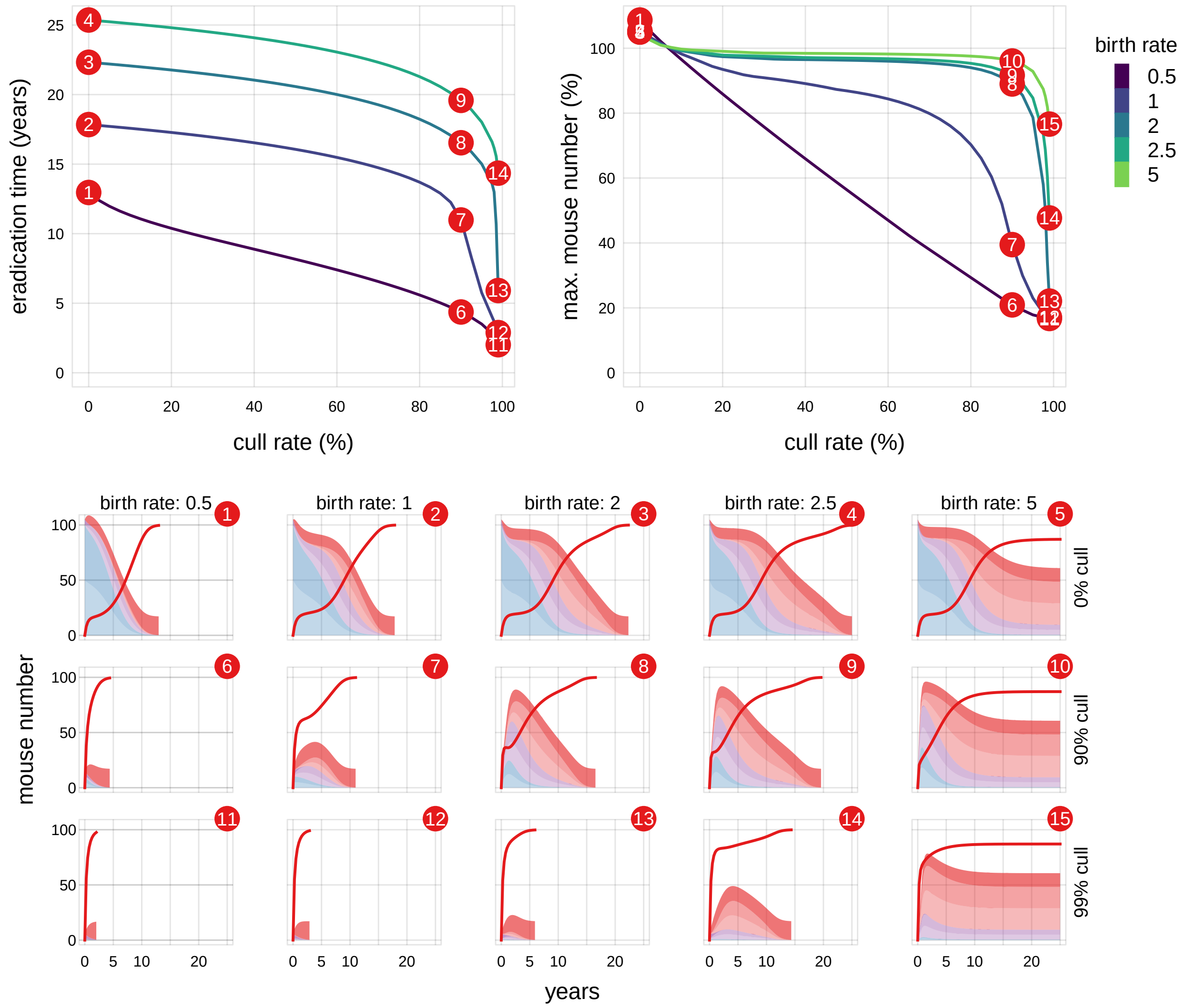


**Figure S4. Cull & release: Efficacy depends on birth rates**
The birth rate $b_0$ (the number of offspring produced per month at $N = 0$; see Fig. 1B) influences how quickly the population recovers following the cull, and therefore affects both the time to eradication and the maximum population size reached during the campaign.
**Top panels.** Eradication time (years, **left**) and maximum population size (% of carrying capacity $K$, **right**) following a cull and subsequent $t$ release campaign. The x axis shows the extent of the cull, and colours indicate five levels of birth rate. Red numbers indicate the scenarios illustrated in the bottom panels. Scenarios that do not result in eradication will not appear in the left panel.
**Bottom panels:** Population dynamics for 15 scenarios illustrating the effects of birth rate and cull extent, with all other parameters held at baseline values (Table 1). Sterile $t/t$ males are released at a rate of 1% per month in all scenarios. Red lines show the proportion of $t/t$ sterile males.

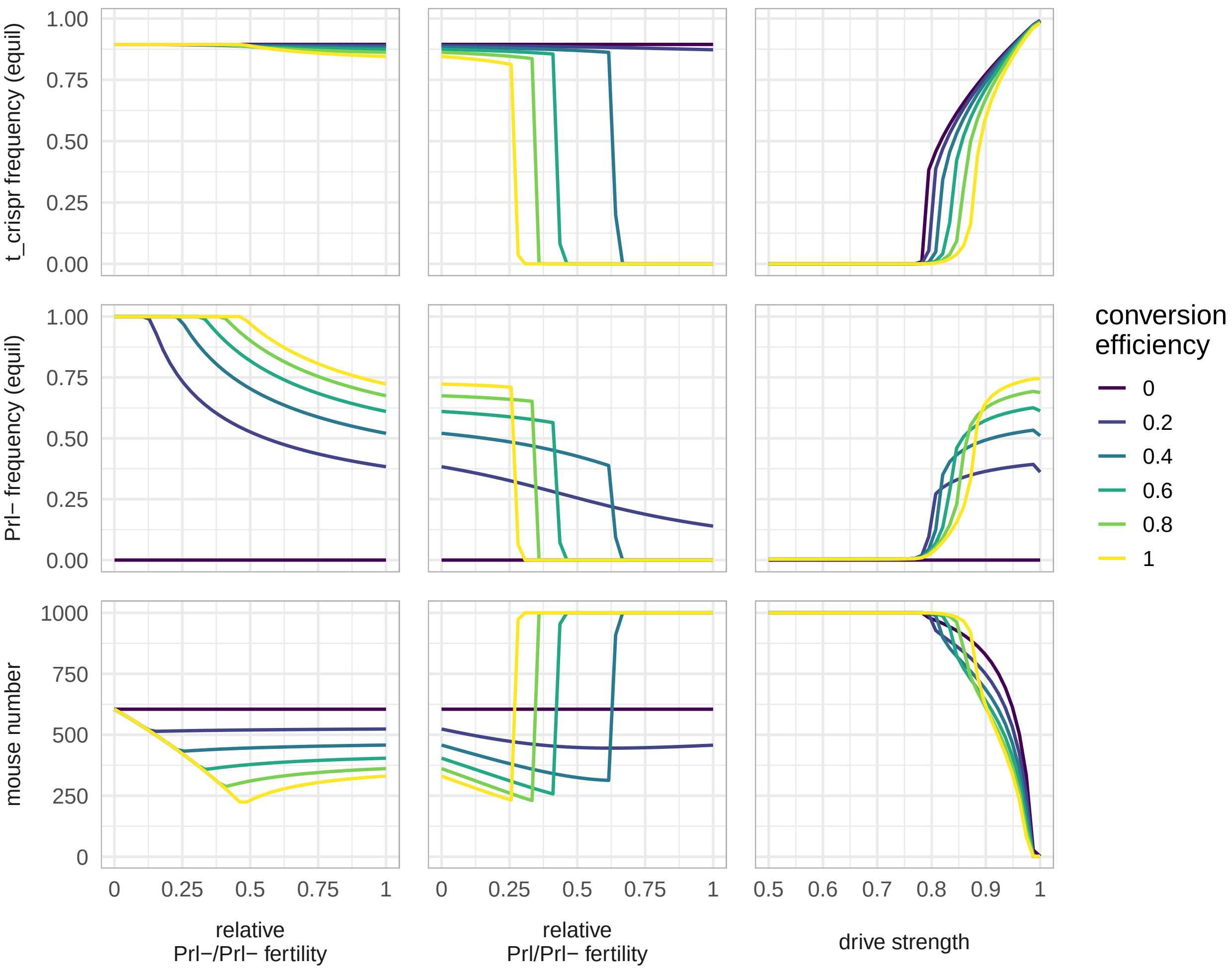


**Figure S5.** $t_{\mathrm{CRISPR}}$**: Effects of *Prl-* fertility load, drive strength, and conversion efficiency**

Population suppression (**bottom row:** final population size) caused by single release of $+/t_{\mathrm{CRISPR}}$ males at 5% reflects the combined effect of $t_{\mathrm{CRISPR}}$ spread (**top row:** equilibrium $t_{\mathrm{CRISPR}}$ frequency) and the resulting loss of function at the *Prl* locus (**middle row:** equilibrium $Prl^{-}$ frequency).

**Left column.** Homozygote $Prl^{-}/Prl^{-}$ fertility $w_{\mathrm{Prl\text{-}/Prl\text{-}}}$ relative to $Prl/Prl$ females whose fertility $w_{\mathrm{Prl/Prl}} = 1$, with heterozygote fertility $w_{\mathrm{Prl/Prl\text{-}}} = 1$. Colors indicate conversion efficiency $c$, remaining parameters are at baseline. **Center column.** Heterozygote fertility $w_{\mathrm{Prl/Prl\text{-}}}$ relative to $Prl/Prl$ females, with $w_{\mathrm{Prl\text{-}/Prl\text{-}}} = 0$ and $c = 0.8$. **Right column.** Drive strength $d$, with baseline fertility $w_{\mathrm{Prl/Prl\text{-}}} = 1$, $w_{\mathrm{Prl\text{-}/Prl\text{-}}} = 0$.

**Interpretation.** Population suppression is *maximal* at *intermediate* fertility costs. When fertility costs are too high, selection against $Prl^{-}$ prevents both $Prl^{-}$ and $t_{\mathrm{CRISPR}}$ from persisting. Higher conversion efficiency $c$ generally increases population suppression, but also causes $t_{\mathrm{CRISPR}}$ and $Prl^{-}$ to be lost at lower fertility-cost thresholds. Drive strength $d$ consistently increases population suppression, with $d < 0.8$ rendering the approach ineffective. Fertility costs may obviously be mechanistically constrained by the choice of fertility locus targeted by $t_{\mathrm{CRISPR}}$.

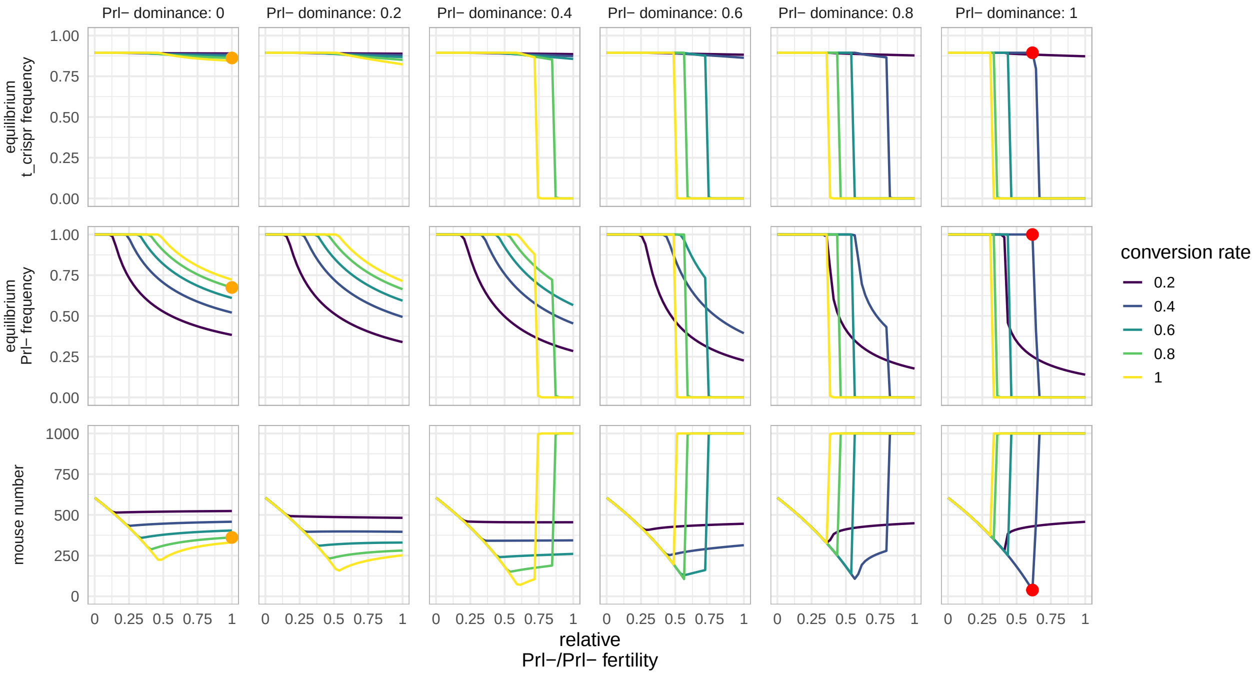


**Figure S6.** $t_{\mathrm{CRISPR}}$**: Fertility load that maximise population suppression**

Although fertility costs at a given locus may be mechanistically constrained and not freely tunable in practice, we here vary them systematically to identify fertility costs that maximise population suppression. Equilibrium frequency of $t_{\mathrm{CRISPR}}$, $Prl^{-}$, and population size are shown as a function of relative homozygous $Prl^{-}/Prl^{-}$ fertility $w_{\text{Prl-/Prl-}}$ (**x axis**), the dominance coefficient for fertility $h_{\text{Prl}}$, where $w_{\text{Prl/Prl-}} = 1 - h_{\text{Prl}}(1 - w_{\text{Prl/Prl-}})$ (**columns**), and conversion efficiency $c$ (**colours**). Remaining parameters are at baseline.

**Interpretation.** Higher conversion efficiencies $c$ increase population suppression, but trigger the complete loss of $Prl^{-}$ and $t_{\mathrm{CRISPR}}$ due to excessive fertility load at lower cost thresholds. Population suppression is generally optimal for intermediate homozygote and heterozygote fertility costs, where $Prl^{-}$ can spread to fixation. Maximum suppression occurs at $w_{\text{Prl-/Prl-}} = 0.61$, $h_{\text{Prl}} = 1$ (thus $w_{\text{Prl/Prl-}} = 0.61$), and $c = 0.4$, where where 3.8% of the population remains (**red dots**). Gierus et al. (3) considered a haplosufficient *Prl* locus, resulting in complete fertility loss in homozygotes but no loss in heterozygotes ($w_{\text{Prl-/Prl-}} = 0$, $h_{\text{Prl}} = 0$, $c = 0.8$, orange dots). Under these conditions, 36% of mice remain at equilibrium, approximately 10 times more than under the optimal fertility profile.

**Caveat.** Optimal fertility costs are close to the threshold beyond which both $t_{\mathrm{CRISPR}}$ and $Prl^{-}$ are lost due to excessive fertility load. It may therefore be difficult to implement reliably if the relevant parameters are uncertain. Other parameters were kept at baseline values (Table 1).

**Table S1. Parameter sensitivity.** Parameter sensitivity of *t/t* male release rate per month required for eradication (in %) and the eradication time at $\mu_{t/t} = 2\%$ (in years) as function of the refinement method and model parameters (corresponding to main text Fig. 3). For each parameter, the following values are shown: **minimum value**, $y_{min}$, for the parameter values considered (including % change compared to the baseline parameter value from Tab. 1); **maximum value**, $y_{max}$, for the parameter range considered; **slope** as $\frac{y_{max}-y_{min}}{x_{y.max}-x_{y.min}}$, where $x_{y.max}$, $x_{y.min}$ denote the parameter values $x$ corresponding to $y_{max}$, $y_{min}$, respectively. The slope thus measure how strongly the outcome variable (e.g. eradication time) changes for each unit change in the parameter value. For example, line 4 tells us the the minimum release rate required for eradication using the *Adonis* method is 0.8% per month, and the eradication time 16.1 years (compared to 1.7% and 16.1 years for the baseline sterile t method). In line 9, we learn that varying turnover at $K$ (range: $m_K = [0.1, 0.6]$) results in changes in required release rates from 0.5 to 0.9%, corresponding to -0.7% change in release rate for each unit increase in $m_K$ (slope). Likewise, eradication time will vary 8.6 to 20.7 years for the same interval of $m_K$ under the *Adonis* method, corresponding to a decrease of 24.2 years per unit increase in $m_k$ (slope).

| Parameter | Method | min release rate (%) | max release rate (%) | slope release rate | min eradication time (years) | max eradication time (years) | slope eradication time |
|---|---|---|---|---|---|---|---|
| baseline | sterile *t* | | **1.7** | | | **25.4** | |
| | 5% trap & release | | **1.0** | | | **17.5** | |
| | 95% cull & release | | **1.6** | | | **18.0** | |
| | *Adonis*, 3-fold | | **0.8** | | | **16.1** | |
| | 80% $t_{CRISPR}$ | | **1.0** | | | **23.3** | |
| turnover at $K$ ($m_K$) | sterile *t* | **1.3** (-25%) | **1.8** (+9%) | **-1.1** | **12.6** (-50%) | **28.3** (+12%) | **-35.1** |
| | 5% trap & release | **0.8** (-24%) | **0.9** (-8%) | **-0.3** | **10.1** (-42%) | **20.2** (+15%) | **-20.1** |
| | 95% cull & release | **1.3** (-20%) | **1.4** (-11%) | **-0.3** | **9.9** (-45%) | **20.0** (+11%) | **-20.3** |
| | *Adonis*, 3-fold | **0.5** (-32%) | **0.9** (+13%) | **-0.7** | **8.6** (-46%) | **20.7** (+29%) | **-24.2** |
| | 80% $t_{CRISPR}$ | **0.5** (-54%) | **1.5** (+51%) | **-2.1** | **12.3** (-47%) | **28.3** (+22%) | **-33.8** |
| birth rate ($b_0$) | sterile *t* | **0.3** (-83%) | **3.0** (+79%) | **0.7** | **17.8** (-30%) | **28.2** (+11%) | **5.8** |
| | 5% trap & release | **0.0** (-99%) | **2.6** (+160%) | **0.7** | **10.7** (-39%) | **25.9** (+48%) | **5.4** |
| | 95% cull & release | **0.1** (-91%) | **2.9** (+82%) | **0.7** | **5.8** (-68%) | **24.0** (+33%) | **9.1** |
| | *Adonis*, 3-fold | **0.1** (-82%) | **1.5** (+98%) | **0.3** | **12.1** (-25%) | **19.8** (+23%) | **1.9** |
| | 80% $t_{CRISPR}$ | **0.2** (-82%) | **2.1** (+111%) | **0.5** | **17.1** (-27%) | **29.3** (+26%) | **3.8** |
| *t/t* competitiveness ($c_{t/t}$) | sterile *t* | **0.6** (-67%) | **2.9** (+70%) | **-0.8** | **19.1** (-25%) | **27.6** (+9%) | **-3.9** |
| | 5% trap & release | **0.1** (-86%) | **2.3** (+123%) | **-0.7** | **14.2** (-19%) | **28.4** (+62%) | **-5.1** |
| | 95% cull & release | **0.4** (-75%) | **2.8** (+74%) | **-0.8** | **12.1** (-33%) | **23.6** (+31%) | **-5.0** |
| | *Adonis*, 3-fold | **0.3** (-65%) | **1.3** (+65%) | **-0.3** | **14.6** (-9%) | **17.6** (+10%) | **-1.0** |
| | 80% $t_{CRISPR}$ | **0.3** (-69%) | **1.7** (+76%) | **-0.5** | **19.8** (-15%) | **28.3** (+22%) | **-2.9** |
| drive strength ($d$) | sterile *t* | **0.0** (-99%) | **4.7** (+178%) | **-31.7** | **16.4** (-35%) | **25.4** (+0%) | **-179.6** |
| | 5% trap & release | **0.0** (-98%) | **4.8** (+370%) | **-24.2** | **12.8** (-27%) | **24.5** (+40%) | **-158.9** |
| | 95% cull & release | **0.0** (-98%) | **5.0** (+216%) | **-29.1** | **10.2** (-43%) | **18.0** (+0%) | **-156.6** |
| | *Adonis*, 3-fold | **0.0** (-98%) | **4.0** (+426%) | **-8.2** | **13.4** (-17%) | **21.6** (+34%) | **-84.0** |
| | 80% $t_{CRISPR}$ | **0.0** (-99%) | **5.0** (+405%) | **-20.3** | **17.7** (-24%) | **28.1** (+21%) | **-142.5** |
| *+/t* sperm competitiveness ($\rho$) | sterile *t* | **1.5** (-10%) | **1.7** (+3%) | **-0.2** | **19.7** (-22%) | **28.8** (+14%) | **-9.2** |
| | 5% trap & release | **0.9** (-12%) | **1.1** (+4%) | **-0.2** | **13.2** (-25%) | **20.3** (+16%) | **-7.1** |
| | 95% cull & release | **1.5** (-4%) | **1.6** (+4%) | **-0.1** | **16.0** (-11%) | **18.7** (+4%) | **-2.8** |
| | *Adonis*, 3-fold | **0.7** (-7%) | **0.8** (+6%) | **-0.1** | **13.6** (-16%) | **17.1** (+6%) | **-3.6** |
| | 80% $t_{CRISPR}$ | **0.7** (-34%) | **1.2** (+25%) | **-0.6** | **16.0** (-31%) | **28.7** (+23%) | **-12.8** |
| sterile mating cost ($t_{miss}$) | sterile *t* | **0.3** (-83%) | **4.5** (+168%) | **-4.7** | **16.6** (-34%) | **27.1** (+7%) | **-14.5** |
| | 5% trap & release | **0.1** (-86%) | **3.6** (+253%) | **-3.8** | **12.6** (-28%) | **25.2** (+44%) | **-15.6** |
| | 95% cull & release | **0.2** (-89%) | **4.4** (+177%) | **-4.7** | **8.9** (-50%) | **19.7** (+9%) | **-15.0** |
| | *Adonis*, 3-fold | **0.1** (-82%) | **2.4** (+213%) | **-2.5** | **9.9** (-39%) | **22.2** (+38%) | **-14.5** |
| | 80% $t_{CRISPR}$ | **0.2** (-76%) | **2.5** (+149%) | **-2.5** | **17.3** (-25%) | **27.1** (+16%) | **-12.0** |
| polyandry ($p$) | sterile *t* | **0.9** (-47%) | **2.7** (+63%) | **1.8** | **15.6** (-38%) | **28.5** (+12%) | **23.4** |
| | 5% trap & release | **0.4** (-61%) | **2.1** (+109%) | **1.7** | **11.3** (-36%) | **28.1** (+60%) | **22.4** |
| | 95% cull & release | **0.9** (-44%) | **2.4** (+51%) | **1.5** | **11.8** (-34%) | **27.8** (+54%) | **21.3** |
| | *Adonis*, 3-fold | **0.4** (-49%) | **1.3** (+71%) | **0.9** | **12.1** (-25%) | **22.9** (+42%) | **10.8** |
| | 80% $t_{CRISPR}$ | **0.4** (-60%) | **3.0** (+203%) | **2.6** | **14.2** (-39%) | **28.8** (+24%) | **24.3** |
| *t/t* mortality ($m_{t/t}$) | sterile *t* | **0.0** (-98%) | **5.1** (+204%) | **2.7** | **6.4** (-75%) | **27.6** (+9%) | **22.2** |
| | 5% trap & release | **0.0** (-98%) | **2.9** (+186%) | **1.5** | **8.8** (-50%) | **27.9** (+60%) | **13.5** |
| | 95% cull & release | **0.1** (-96%) | **4.9** (+209%) | **2.6** | **4.8** (-73%) | **28.2** (+56%) | **22.4** |
| | *Adonis*, 3-fold | **0.0** (-97%) | **2.3** (+202%) | **1.2** | **5.2** (-68%) | **29.8** (+86%) | **14.4** |
| | 80% $t_{CRISPR}$ | **0.0** (-98%) | **3.0** (+203%) | **1.6** | **6.4** (-72%) | **29.5** (+27%) | **18.7** |
